# Hierarchical Stem Cell Topography Splits Growth and Homeostatic Functions in the Fish Gill

**DOI:** 10.1101/484139

**Authors:** Julian Stolper, Elizabeth Ambrosio, Diana-P Danciu, David A. Elliot, Kiyoshi Naruse, Anna Marciniak Czochra, Lazaro Centanin

**Affiliations:** Centre for Organismal Studies, Heidelberg University, 69120 Heidelberg, Germany; Murdoch Children’s Research Institute, Royal Children’s Hospital, Parkville, VIC, 3052, Australia; Institute of Applied Mathematics, Interdisciplinary Center for Scientific Computing (IWR) and BIOQUANT Center, Heidelberg University, 69120 Heidelberg, Germany; Laboratory of Bioresources, National Institute for Basic Biology, National Institutes of Natural Sciences, Okazaki, Aichi 444-8585, Japan; Current address: Danish Stem Cell Center (DanStem), University of Copenhagen, 2200 Copenhagen N, Denmark

## Abstract

While lower vertebrates contain adult stem cells (aSCs) that maintain homeostasis and drive un-exhaustive organismal growth, mammalian aSCs display mainly the homeostatic function. Understanding aSC-driven growth is of paramount importance to promote organ regeneration and prevent tumor formation in mammals. Here we present a clonal approach to address common or dedicated populations of aSCs for homeostasis and growth. Our functional assays on medaka gills demonstrate the existence of separate homeostatic and growth aSCs, which are clonal but differ in their topology. While homeostatic aSCs are fixed, embedded in the tissue, growth aSCs locate at the expanding peripheral zone. Modifications in tissue architecture can convert the homeostatic zone into a growth zone, indicating a leading role for the physical niche defining stem cell output. We hypothesize that physical niches are main players to restrict aSCs to a homeostatic function in animals with a fixed adult size.

## Introduction

Higher vertebrates acquire a definitive body size around the time of their sexual maturation. Although many adult stem cells (aSCs) remain active and keep producing new cells afterwards, they mainly replace cells that are lost on a daily basis. On the other hand, lower vertebrates like fish keep increasing their size even during adulthood due to the capacity of aSCs to drive growth in parallel to maintaining organ homeostasis. The basis for the different outputs between aSCs in lower and higher vertebrates is still not fully understood. It has been reported, however, that in pathological conditions mammalian aSCs exhibit the ability to drive growth, as best represented by cancer stem cells (CSCs) (Batlle & Clevers, 2017; Nassar & Blanpain, 2016; Clevers, 2011; Suvà *et al*, 2014; Quintana *et al*, 2008; Barker *et al*, 2009; Schepers *et al*, 2012; Boumahdi *et al*, 2014).

Since stem cells in fish maintain homeostasis and drive post-embryonic growth in a highly controlled manner, the system permits identifying similarities and differences in case both functions are performed by dedicated populations, or identifying conditions for homeostatic and growth outputs in case of a common stem cell. There are several genetic tools and techniques to explore aSCs in fish, and an abundant literature covering different aspects of their biology in various organs and also during regeneration paradigms (Gupta & Poss, 2012; Knopf *et al*, 2011; Tu & Johnson, 2011; Kizil *et al*, 2012; Kyritsis *et al*, 2012; Pan *et al*, 2013; Centanin *et al*, 2014; Jungke *et al*, 2015; Henninger *et al*, 2017; Singh *et al*, 2017; McKenna *et al*, 2016; Aghaallaei *et al*, 2016). Despite all these major advances, we still do not understand whether the same pool of stem cells is responsible for driving both growth and homeostatic replacement, or if alternatively, each task is performed by dedicated aSCs.

We decided to address this question using the medaka gill, which works as a respiratory, sensory and osmoregulatory organ in most teleost fish. Gills are permanently exposed to circulating water and therefore have a high turnover rate (Chrétien & Pisam, 1986). Additionally, their growth pace must guarantee oxygen supply to meet the energetic demands of a growing organismal size. Moving from the highest-level structure to the smallest, gills are organised in four pairs of branchial arches, a number which remains constant through the fish’s life. Each brachial arch consists of two rows of an ever-increasing number of filaments that are added life-long at both extremes (Figure 1A). Primary filaments have a core from which secondary filaments, or lamellae, protrude. The lamellae are the respiratory unit of the organ, and new lamellae are continually produced within each filament (Wilson & Laurent, 2002). Bigger fish therefore display more filaments that are longer than those of smaller fish, and there is a direct correlation of filament length and number and the body size of the fish (Wilson & Laurent, 2002).

**Figure 1.**
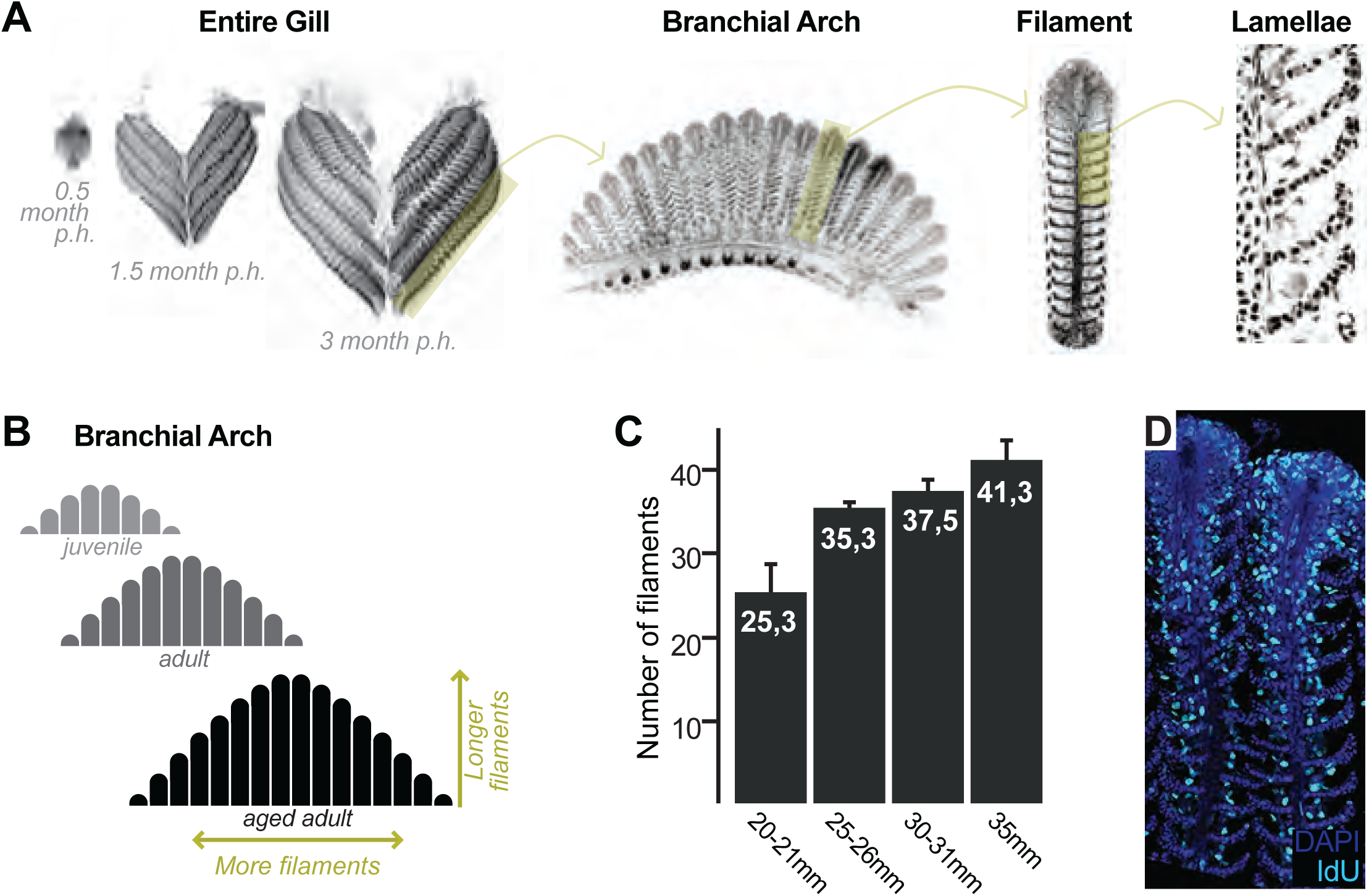
Growth and Homeostasis in the Medaka Gill. (**A**) Enucleated entire gills of medaka at different post-embryonic times show that organ size increases during post-embryonic growth (***left***). A gill contains 4 pairs of branchial arches (***middle left***) that display numerous filaments (***middle right***). Filaments are composed of lamella (***right***), where gas exchange occurs. (**B**) Scheme depicting that branchial arches grow by increasing the number of filaments, and filaments grow by increasing its length. (**C**) The number of filaments per branchial arch is higher in bigger fish - x axis represents fish length, and y axis the number of filaments in the second right branchial arch. (**D**) IdU incorporation in the adult gill reflects proliferating cells all along the longitudinal axis of a filament.

Besides being the respiratory organ of fish, the gill has additional functions as a sensory and osmoregulatory organ (Sundin & Nilsson, 2002; Wilson & Laurent, 2002; Jonz & Nurse, 2005; Hockman *et al*, 2017). It contains oxygen sensing cells (Jonz *et al*, 2004), similar to those found in the mammalian carotid body although with a different lineage history (Hockman *et al*, 2017), and mitochondrial rich cells (MRCs) (Wilson & Laurent, 2002) that regulate ion uptake and excretion and are identified by a distinctive Na+, K+, ATPase activity. Other cell types include pavement cells (respiratory cells of the gills), pillar cells (structural support for lamellae), globe cells (mucous secretory cells), chondrocytes (skeleton of the filaments) and vascular cells. All these cell types must be permanently produced in a coordinated manner during the post-embryonic life of fish. The gill constitutes, therefore, an organ that allows addressing adult stem cells during the addition and homeostatic replacement of numerous, diverse cell types.

Bona fide stem cells can only be identified and characterised by following their offspring for long periods to prove self-renewal, the defining feature of stem cells (Clevers & Watt, 2018). In this study, we use a lineage analysis approach that revealed growth and homeostatic stem cells in the medaka gill. We found that gill stem cells are fate restricted, and identified at least four different lineages along each filament. By generating clones at different stages, we show that these four lineages are generated early in embryogenesis, previous to the formation of the gill. Our results also indicate that growth and homeostatic aSCs locate to different regions along the gill filaments and the branchial arches. Homeostatic stem cells have a fixed position embedded in the tissue, and generate cells that move away to be integrated in an already functional unit, similarly to mammalian aSCs in the intestinal crypt (Barker *et al*, 2008). Growth stem cells, on the other hand, locate to the growing edge of filaments and are moved as filaments grow, resembling the activity of plant growth stem cells at the apical meristems (Greb & Lohmann, 2016). We have also found that the homeostatic aSCs can turn into growth aSCs when the apical part of a filament is ablated, revealing that the activity of a stem cell is highly plastic and depends on the local environment. Our data reveal a topological difference between growth and homeostatic stem cells, that has similar functional consequences in diverse stem cell systems.

## Results

### Medaka Gills Contain Homeostatic and Growth Stem Cells

The fish gill displays a significant post-embryonic expansion that reflects the activity of growth stem cells and a fast turnover rate that indicates the presence of homeostatic cells. Gills massively increment their size during medaka post-embryonic life (Figure 1A, left), where growth happens along two orthogonal axes. One axis represents increase in length of each filament, and the other, the iterative addition of new filaments to a branchial arch. This way, branchial arches of an adult fish contain more filaments, which are also longer, than those of juveniles. Branchial arches in medaka continue to expand along these two axes well after sexual maturation (Figure 1B, C). Gills from teleost fish are exposed to the surrounding water and experience a fast turnover rate. When adult medaka are incubated with IdU for 48 h, their gill filaments display a strong signal from the base to the top (Figure 1D), which indicates the presence of mitotically active cells all along the filament’s longitudinal axis. These observations position medaka gills as an ideal system to explore the presence of growth and homeostatic stem cells within the same organ and address their similarities and differences.

### Growth Stem Cells Locate to Both Growing Edges of Each Branchial Arch

We first focussed on identifying growth stem cells, by combining experimental data on clonal progression with a mathematical approach to quantify the expected behavior for stem cell- and progenitor-mediated growth. Experimentally, clones were generated using the Gaudí toolkit, which consists of transgenic lines bearing floxed fluorescent reporter cassettes (Gaudí^*RSG*^ or Gaudí^*BBW2.1*^) and allows inducing either the expression or the activity of the Cre recombinase (Gaudí^*Hsp70A.CRE*^ or Gaudí^*Ubiq.iCRE*^, respectively). The Gaudí toolkit has already been extensively used for lineage analyses in medaka (Centanin *et al*, 2014; Reinhardt *et al*, 2015; Lust *et al*, 2016; Aghaallaei *et al*, 2016; Seleit *et al*, 2017). Clones are generated by applying subtle heat-shock treatments (when Gaudí^*Hsp70A.CRE*^ is used) or low doses of tamoxifen (when Gaudí^*Ubiq.iCRE*^ is used) to double transgenic animals, which results in a sparse labelling of different cells along the fish body, transmitting the label to their offspring.

The length of filaments increases from peripheral to central positions (Figure 1A, 2A), regardless of the total number of filaments per branchial arch (Leguen, 2017). This particular arrangement suggests that the oldest and therefore longest filaments, of embryonic origin, locate to the centre of a branchial arch, while the new filaments are incorporated at the peripheral extremes either by stem cells (permanent) or progenitors (exhaustive). Conceptually, the latter two scenarios would lead to different lineage outputs. If filaments were formed from progenitor cells that are already present at the time of labelling, we would anticipate that the post-embryonic - peripheral - domain of adult branchial arches should contain both labelled and unlabelled filaments (Figure 2A, bottom left). Alternatively, if post-embryonic filaments were generated by a *bona fide*, self-renewing stem cells, the periphery of adult branchial arches should be homogeneous in its labelling status, containing either labelled or non-labelled stretches of clonal filaments (Figure 2A, bottom right). When we analysed adult Gaudí^*Ubiq.iCRE*^ Gaudí^*RSG*^ transgenic fish that had been induced for sparse recombination at old embryonic stages (9 dpf.), we observed that post-embryonic filaments at the extreme of branchial arches were grouped in either labelled or non-labelled stretches (Figure 2B, C, asterisks for labelled stretches and arrowheads for embryonic filaments) suggesting that they were generated by bona-fide stem cells.

**Figure 2.**
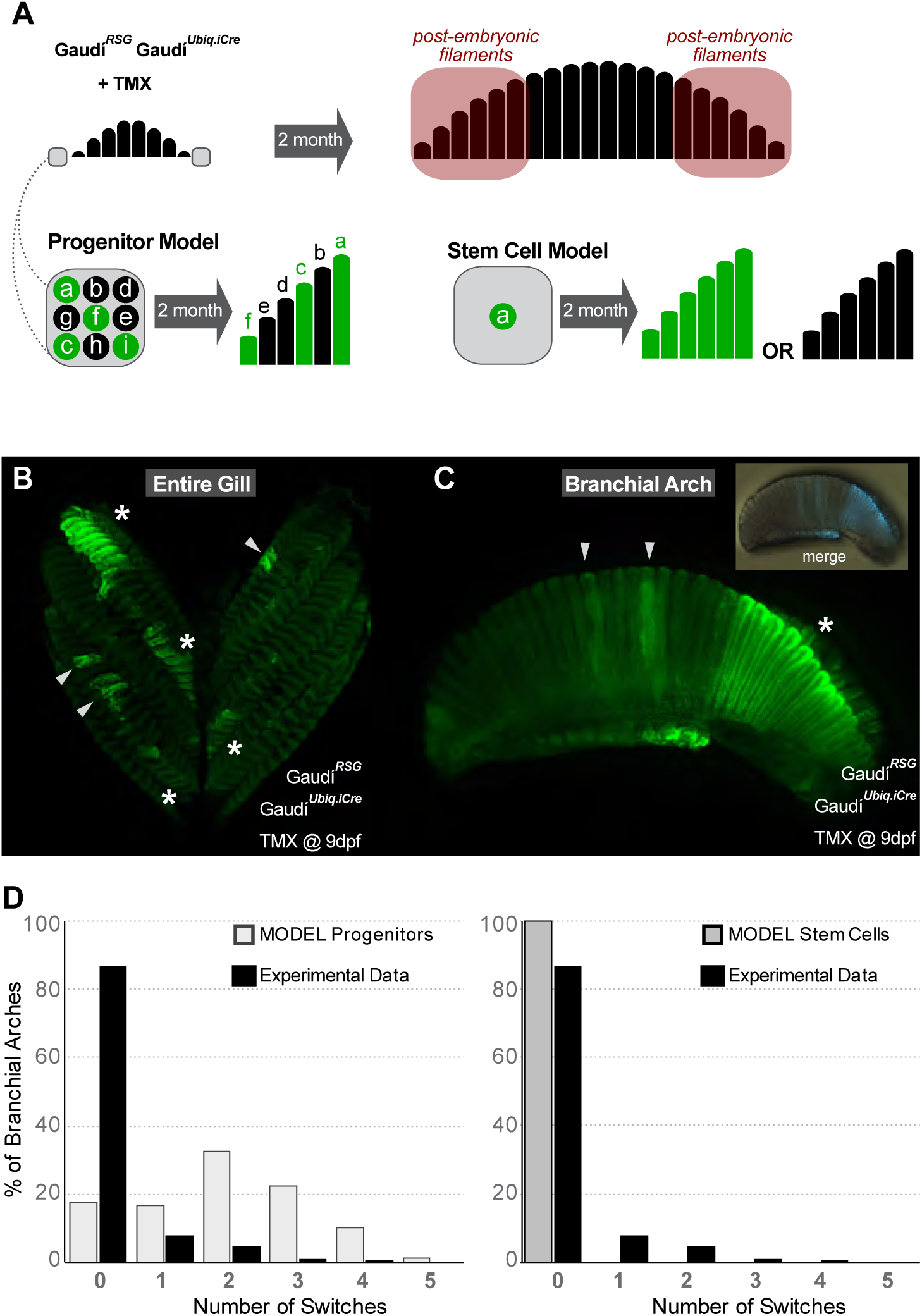
Gill Stem Cells Located at the Periphery of Branchial Arches Generate More Filaments Life-Long. (**A**) Scheme showing the expected outcome assuming a progenitor (***left bottom***) or a stem cell (***right bottom***) model. (**B**) Entire gill from a double transgenic Gaudí^*Ubiq.iCre*^ Gaudí^*RSG*^ fish 2 month after induction with TMX. (**C**) Branchial arch from a double transgenic Gaudí^*Ubiq.iCre*^ Gaudí^*RSG*^ fish two months after induction with TMX. Arrowheads in B and C indicate recombined embryonic filaments located at the centre of branchial arches, and asterisks indicate stretches of peripheral filaments with the same recombination status. (**D**) Graphs showing the distribution of switches in stretches of the 6 most peripheral filaments. The graphs show a comparison of the experimental data (black) to the expected distribution according to a progenitor model (light gray, left) and to a stem cell model (gray, right).

Our experimental data were then compared to the outcome of a computational model accounting for different scenarios for progenitor and stem cell mediated growth. The analysis was focussed on the six most peripheral filaments of adult branchial arches (See M&M for details on filament numbers and how labelling efficiency was calculated). For each scenario, we employed stochastic simulations assigning "0"to a non-labelled filament and "1" to a labelled filament and computing the number of switches in the labelled status of two consecutive filaments, i.e. the number of transitions from “0-to-1” and from “1-to-0” (Supplementary Tables 1 and 2) (1,000 simulations on 5,000 randomly generated stretches for each experimental gill analysed, see M&M). Assuming a labelling efficiency of 50%, a progenitor-based model results in a normal distribution of switches while a stem-cell-based model shows no switches among consecutive filaments, i.e. contains only filaments that have a value of either 0 or 1 (see Figure 2D for the number of switches for each model with labelling efficiencies estimated from experiments). We have quantified both peripheral extremes of hundreds of experimental branchial arches (N >300 6-*filament* stretches, N=22 independent gills) (Supplementary Table 3) and compared each individual branchial arch to the simulation results of the two models. For every gill analysed, the *stem cell* model explained the experimental data better than the *progenitor cell* model (Supplementary Table 4). Altogether, our data revealed the existence of growth stem cells at the peripheral extremes of branchial arches, which generate new filaments during the post-embryonic life in medaka.

### Growth Stem Cells Locate to the Growing Edge of Each Filament

The massive post-embryonic growth of teleost gills occurs by increasing the number but also the length of filaments. Previous data on stationary samples suggest that filaments grow from their tip (Morgan, 1974), and we followed two complementary dynamic approaches to characterise stem cells during filament growth. First, we exploited the high rate of cellular turnover previously observed by a pulse of IdU (Figure 1D) which labels mitotic cells all along the filament. We reasoned that during a chase period, cells that divide repeatedly — as expected for stem cells driving growth — would dilute their IdU content with every cell division, as previously reported for other fish tissues (Centanin *et al*, 2011). Therefore, the chase period reveals a region in the filament with a decreased signal for IdU that may in turn indicate where new cells are being added (Figure 3A illustrates the different scenarios). Indeed, all filaments analysed contained a region deprived of IdU at the most distal tip (Figure 3B), what stays in agreement with the previous assumptions. Complementary, we performed a clonal analysis by inducing sparse recombination using Gaudí transgenic fish. To reveal the localisation of growing clones, Gaudí^*Ubiq.iCRE*^ Gaudí^*RSG*^ fish were induced for recombination at 3 weeks post fertilisation and grown for 2 months after tamoxifen treatment. We observed that clones at the proximal and middle part of the filament were small and restricted to one lamellae, while the clones at the distal part contained hundreds of cells suggesting that they were generated by growth stem cells (Figure 3C, D).

**Figure 3.**
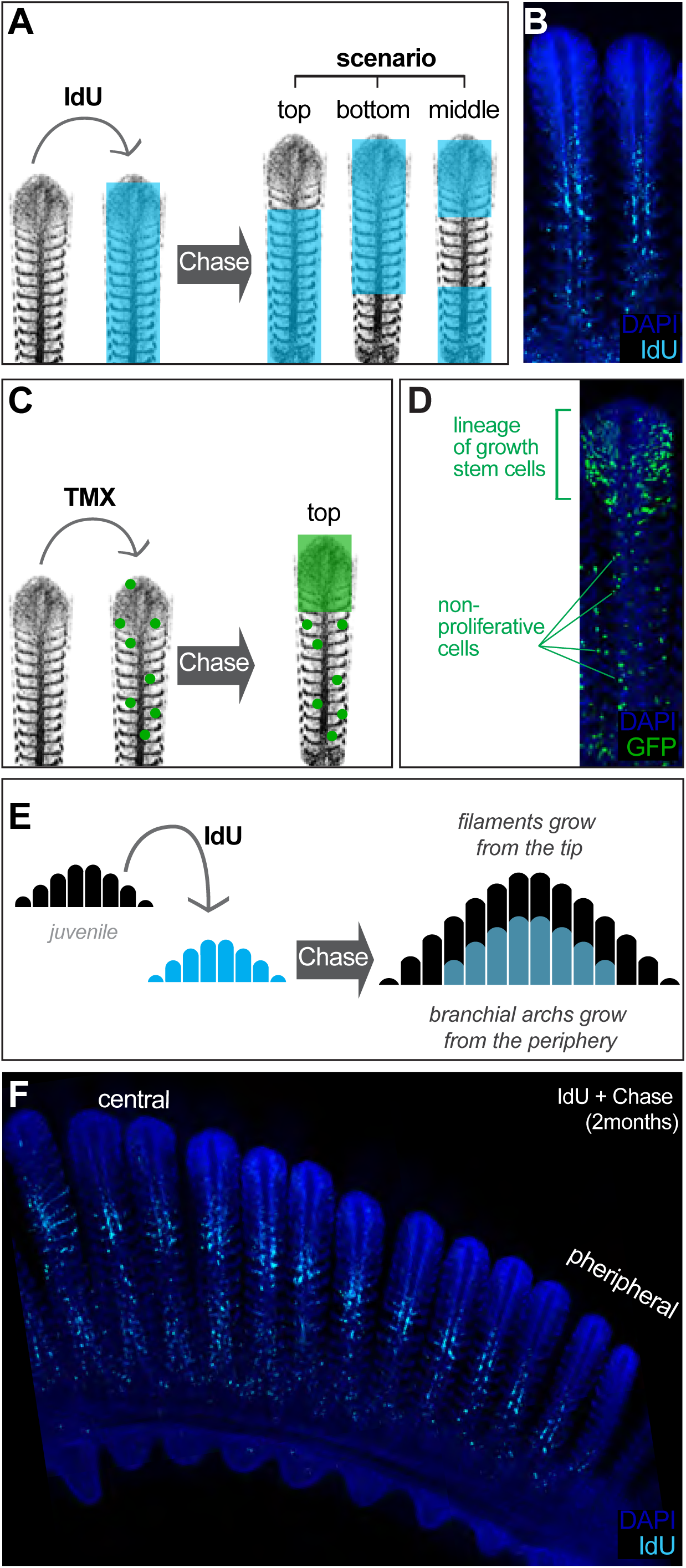
Filament Growth Stem Cells are Located at the Apical Tip. (**A**) Scheme showing the expected outcome of IdU *pulse & chase* experiments depending on the location of growth stem cells. (**B**) IdU *pulse & chase* experiment shows the apical region devoted of signal, indicating these cells were generated after the IdU pulse. (**C**) Scheme showing the expected outcome of a filament in which growth stem cells were labelled. (**D**) A filament from a double transgenic Gaudí^*Ubiq.iCre*^ Gaudí^*RSG*^ fish one month after induction with TMX shows an expanding clone in the apical region, indicating a high proliferative activity compared to clones located at other coordinates along the longitudinal axis. (**E, F**). Scheme (E) and data (F) showing an IdU *pulse & chase* experiment on branchial arches. The apical part of each filament and the more peripheral filaments are devoted of signal revealing the stereotypic growth of branchial arches.

Analysis of pulse-chase IdU experiments in entire branchial arches also suggested that the fraction of IdU labelled cells decreased from central to peripheral filaments. While the most central filaments contain IdU positive cells in roughly 80% of their length, filaments close to the periphery contain just few IdU cells at the basal part or even no IdU cell at all, indicating that they were produced after IdU administration. Macroscopically, IdU label had a shape of a smaller-sized branchial arch nested within a non-labelled, bigger branchial arch (Figure 3E, F). Interestingly, while we observed that the central filaments showed a longer basal signal that becomes shorter in more peripheral filaments, the upper non-labelled fraction seemed rather stable along central-to-periphery axis of the branchial arch (Figure 3F). This suggested that individual filaments had grown at comparable rates during the chase phase, highlighting a coordination among the stem cells that sustained length growth in each filament. Taken together, IdU experiments revealed growth of filaments starting from their most distal extreme, and clonal analysis indicated location of the growth stem cells at the growing tip of each filament.

### Growth Stem Cells are Fate Restricted

Gill filaments contain different cell types distributed along their longitudinal axis (Laurent, 1984; Sundin & Nilsson, 2002; Wilson & Laurent, 2002). Having revealed growth stem cells at the tip of each filament, we explored whether different cell types had a dedicated or a common stem cell during post-embryonic growth. Previous experiments in zebrafish on labelling cell populations at early embryonic stages revealed that neuro-endocrine cells (NECs) are derived from the endoderm (Hockman *et al*, 2017), while pillar cells have a neural crest origin (Mongera *et al*, 2013). We followed a holistic approach to address the potency of gill stem cells once the organ is formed, by using inducible ubiquitous drivers to potentially label all possible lineages within a gill filament. We induced sparse recombination at 8 dpf. in Gaudí^*Ubiq.iCRE*^ Gaudí^*RSG*^ double transgenic fish and grew them to adulthood. We selected gills with EGFP positive clones (Figure 4A), and imaged branchial arches and gill filaments with cellular resolution (Figure 4B-F). Our analysis revealed the presence of four different recombination patterns illustrating the lineage of different types of growth stem cells (Figure 4C-F, patterns 1 to 4). Moreover, this lineage analysis approach showed that growth stem cells at the tip of gill filaments are indeed fate restricted, and hence, the most apical domain of a filament hosts different growth stem cells with complementary potential.

**Figure 4.**
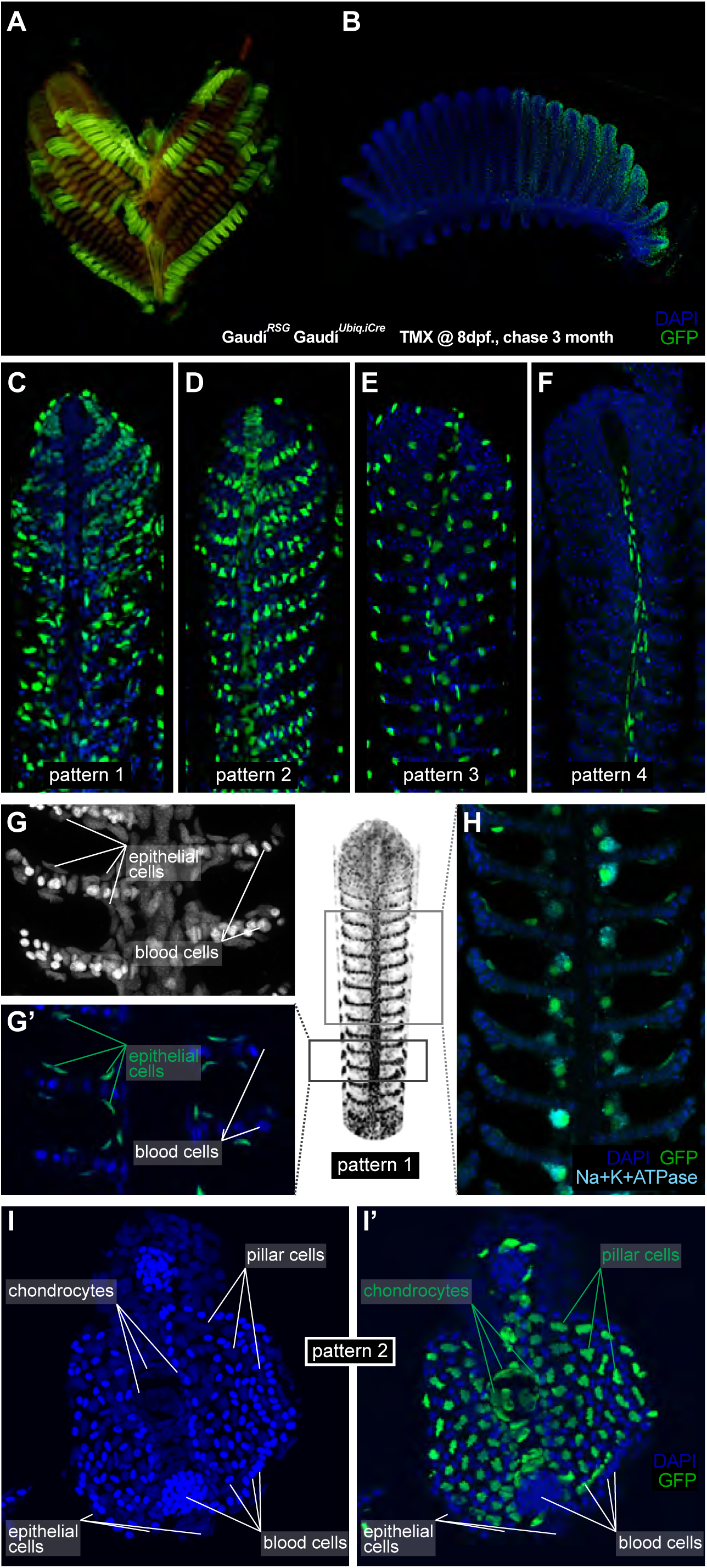
Filament Growth Stem Cells are Fate Restricted. (**A-B**) A gill (A) and a branchial arch (B) from a double transgenic Gaudí^*Ubiq.iCre*^ Gaudí^*RSG*^ fish two month after induction with TMX. (**C-F**) Confocal images from filaments in A, B, stained for EGFP and DAPI to reveal the cellular composition of different clones. Four different recombination patterns were identified. (**G, G’**) A detailed view of Pattern 1(C) show recombined epithelial cells covering each lamella. (**H**) Co-staining with an anti-Na^+^K^+^ATP-ase antibody confirms that MRC cells are clonal to other epithelial cells in the filament. (**I, I’**) Cross-section of a filament that displays Pattern 2 (**D**). DAPI staining allows identifying blood cells (strong signal, small round nuclei), pillar cells (weaker signal, star-shaped nuclei), and chondrocytes (elongated nuclei at the central core of the filament) (I). The lineage tracker EGFP reveals that chondrocytes and pillar cells are clonal along a filament (I’).

Noticeable, recombined filaments displayed the same lineage patterns spanning from their base, i.e. juvenile domain, to their tip, i.e. adult domain, (Figure 4C-F) (N > 200 recombined filaments) indicating that growth stem cells maintain both their activity and their potency during a life-time. A detailed description of the different cell types included in each lineage largely exceeds the scope of this study. Broadly speaking, labelled cells in pattern 1 (Figure 4 C, G-H) are epithelial cells covering the lamellae and the interlamellar space, including MRC cells as revealed by expression of the Na+/K+ ATPase (Figure 4H). Pattern 3 and 4 display a reduced number of labelled cells, sparsely distributed along the filament (pattern 3) or surrounding the gill ray (pattern 4) (Supplementary Movies 1 and 2, respectively). Pattern 2 consists of labelled pillar cells and chondrocytes of the gill ray (Figure 4 I, I’, and reconstructions in Supplementary Movie 3), both easily distinguishable by their location and unique nuclear morphology. Both cell types were previously reported as neural crest derivatives (Mongera *et al*, 2013), and our results demonstrate that they are produced by a common stem cell in every filament during the post-embryonic growth of medaka.

We revealed in the previous sections that growth stem cells at the periphery of branchial arches (*brarch*SCs) generate new filaments, and we showed that each filament contains, in turn, growth stem cells (*filam*SCs) of different fates. To address whether the fate of *filam*SCs is acquired when filaments are formed or set up already in *br-arch*SCs and maintained life-long, we exploited the stretches of labelled — and therefore clonal — filaments observed at the periphery of branchial arches in adult Gaudí^RSG^ Gaudí^*Ubiq.iCRE*^ fish induced for recombination during embryogenesis (Figure 2B, 4A, B). We reasoned that if a labelled *br-arch*SC is fate restricted, the consecutive filaments formed from it should display an identical recombination pattern, since *filam*SCs would have inherited the same fate-restriction from their common *br-arch*SC. Alternatively, if *filam*SCs would acquire the fate-restriction when each filament is formed, then a stretch of clonal filaments should display different recombination patterns, based on the independent fate acquisition at the onset of filament formation (schemes in Figure 5A). We have focussed on 153 branchial arch extremes that started with a labelled filament (N= 83 for *rec. pattern 1*, N= 44 for *rec. pattern 2*, N= 22 for *rec. pattern 3* and N= 4 for *rec. pattern 4*), and 97.4% were followed by a filament with the same recombination pattern (Supplementary Table 4). Moreover, 81.7% of stretches maintained the same recombination pattern for 6 or more filaments, indicating that the labelled cell-of-origin for post-embryonic filaments was already fate restricted. Altogether, our data revealed that a branchial arch contains fate restricted growth *br-arch*SCs at its peripheral extremes that produce growth *filam*SCs stem cells with the same fate-restriction.

**Figure 5.**
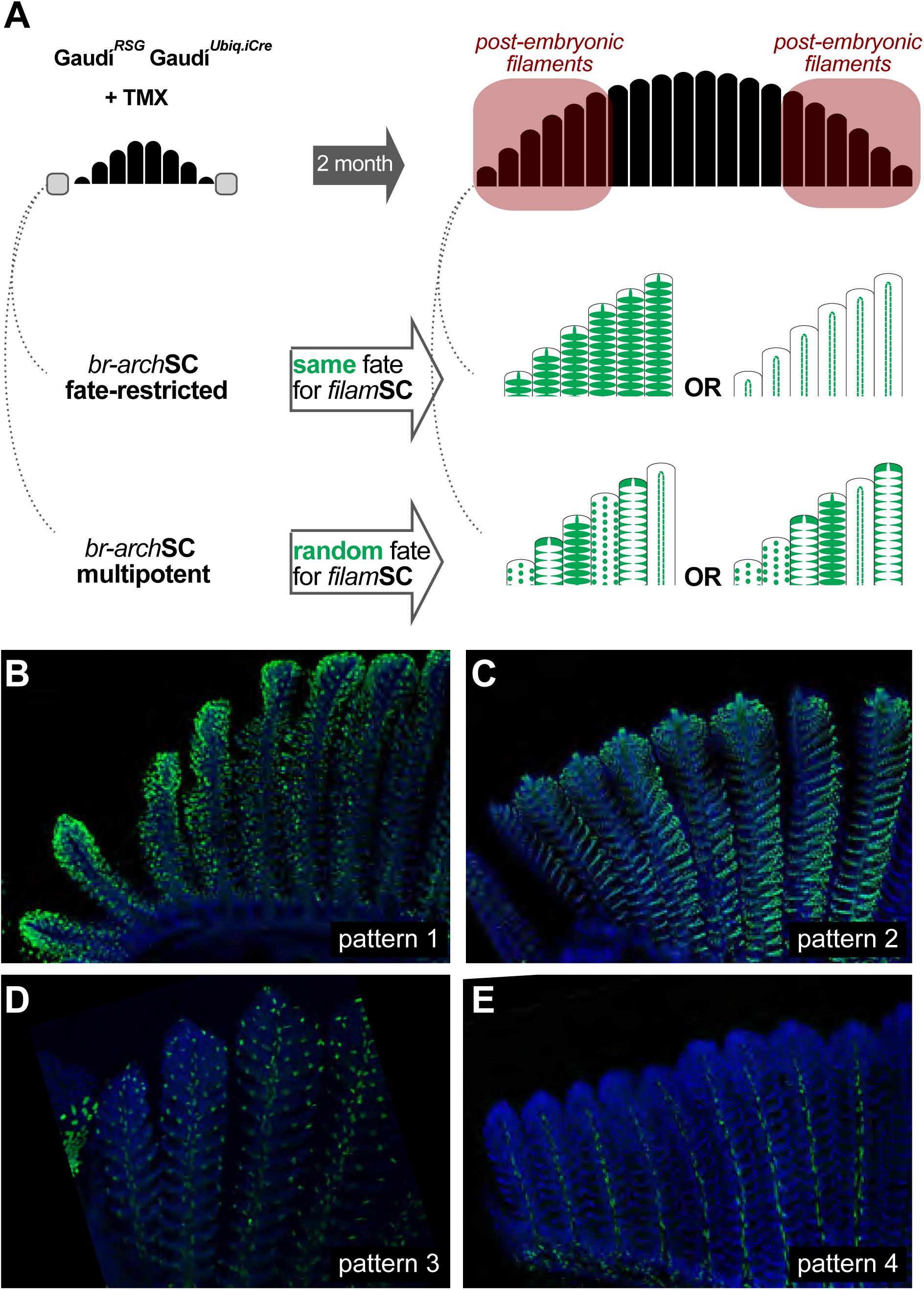
Branchial Arch Stem Cells are Fate Restricted. (**A**) Scheme showing the expected outcome assuming that *br-arch*SCs are fate restricted (***middle***) or multi-potent (***bottom***). The recombination pattern of consecutive filaments would be identical if generated by fate restricted *br-arch*SCs, and non-identical if derived from a multipotent *br-arch*SC. (**B-E**) Confocal images show an identical recombination pattern for peripheral filaments for Pattern 1 (B), Pattern 2 (C), Pattern 3 (D) and Pattern 4 (E).

### Homeostatic Stem Cells Locate to the Base of Each Lamella

Branchial arches grow during post-embryonic life by adding more filaments (Figure 1A-C), and filaments grow in length by adding more lamellae (Figure 1A, Figure 6A). Noticeable, the length of consecutive lamellae does not increase with time along a filament (Figure 6B), resulting in basal and apical lamellae having comparable sizes (basal: 35,72+/-1,93 um and apical: 34,38+/-4,04 um N=6 lamellae of each). This also holds true when comparing the length of lamellae from long (central, embryonic) and short (peripheral, post-embryonic) filaments, and comparing lamellae from medaka of different body length. Lamellae therefore maintain their size despite containing proliferative cells (Laurent, 1984; Laurent *et al*, 1994), a scenario that resembles most mammalian stem cell systems in adults, such as the intestinal crypt or the hair follicle. Previous studies have reported mitotic figures along the filament core in histological sections of various teleost fish. To address the presence and location of proliferating cells in the lamellae of medaka, we performed shorter IdU pulses (12h) and observed that most lamellae contained positive cells at the proximal extreme (Figure 6C), adjacent to the central blood vessels and the gill ray.

**Figure 6.**
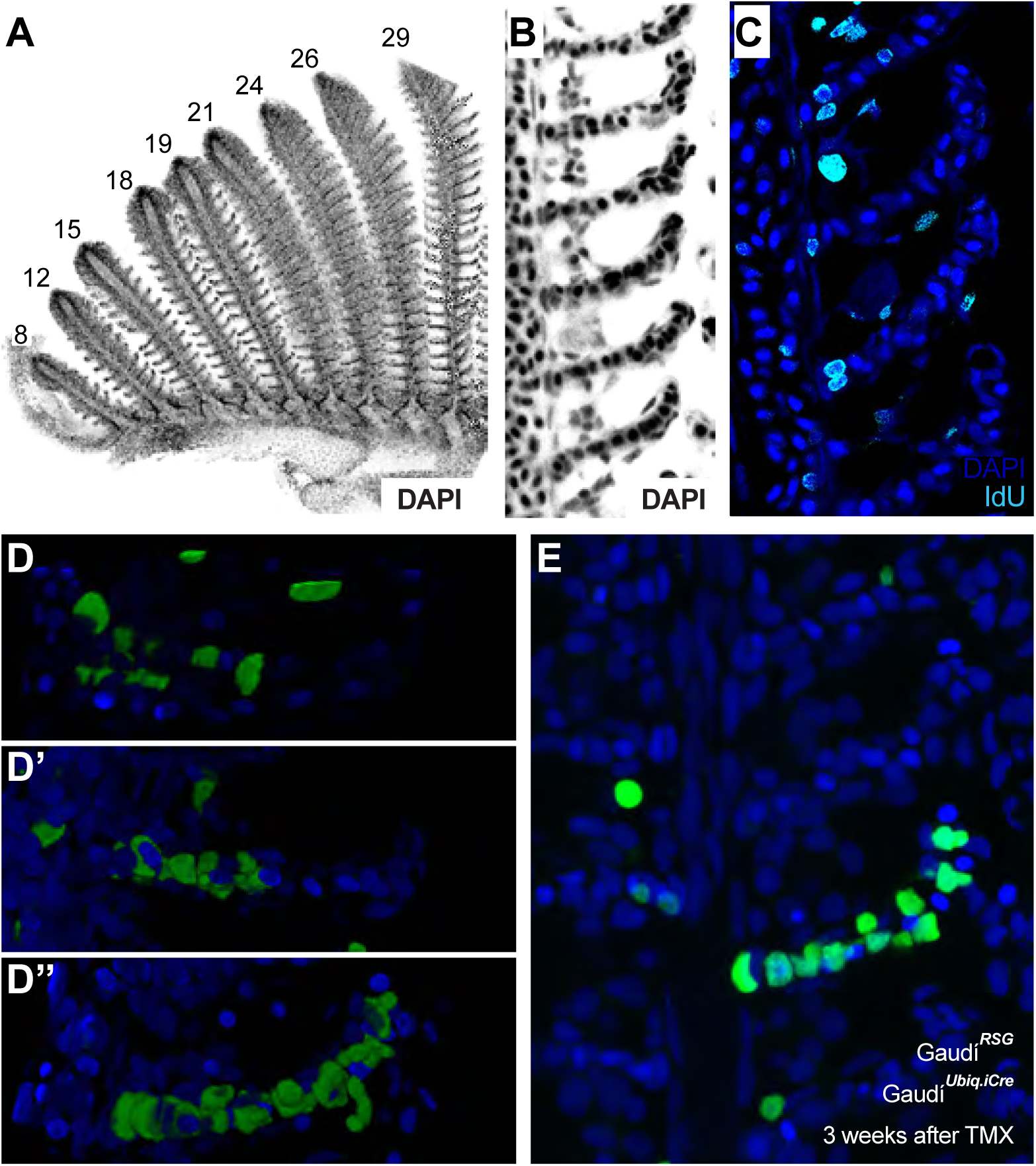
Homeostatic Stem Cells Locate to the Base of Each Lamella. (**A**) DAPI image of peripheral filaments indicating the increasing number of lamellae per filament. (**B**) DAPI image of consecutive lamellae along a filament reveals that lamellae do not increase their size. (**C**) IdU pulse reveals proliferative cells at the base of the lamellae. (**D-E**) EGFP cells indicating clonal progression of clones in double transgenic Gaudí^*Ubiq.iCre*^ Gaudí^*RSG*^ fish one month after induction with TMX during adulthood. Clones of pillar cells progress from the base to the distal part of a lamellae (D’’, E).

We next performed a lineage analysis of gill stem cells during homeostasis, focussing on the lamellae since they constitute naturally-occurring physical compartments that facilitate the analysis of clonal progression. We used double transgenic Gaudí^*Ubiq.iCre*^ Gaudí^*RSG*^ adults that were grown for 3 additional weeks after clonal labelling, and focussed on those containing only a few recombined lamellae per branchial arch (labelling efficiency less than 0.5%). A detailed analysis on lamellae located far away from the filaments’ growing tip revealed clones of labelled cells spanning from the proximal to the distal extreme of the lamella (Figure 6D, E). The clones ranged from a few pillar cells (Figure 6D, D’) to most pillar cells in the lamella (Figure D’’, E and Supplementary Movie 4). This dataset reflects the activity of stem cells contributing to a structure that does not increase in size but renews the cells within — i.e., homeostatic stem cells. Our results therefore indicate the presence of homeostatic pillar stem cells at the base of each lamella in medaka gills.

### The Homeostatic Domain Can Restore Filament Growth

Our lineage analysis revealed distinct locations for both growth and homeostatic stem cells along gill filaments. The growth domain of filaments is always at the top, while the homeostatic domain extends along the longitudinal axis (Figure 7A). Our lineage analysis also revealed that growth and homeostatic stem cells are clonal, since all homeostatic stem cells within a lineage are labelled when a filament has the corresponding labelled growth *filam*SC (Figure 4C-F). We then wondered about their different behaviour; while growth stem cells are displaced by the progeny they generate, homeostatic stem cells maintain their position while pushing their progeny away. These different locations along the filament might constitute dissimilar physical niches. It has indeed been shown in other teleost fish that the growing edge where growth stem cells host is subjected to less spatial restriction than the gill ray niche {Morgan:1974fi}. On the other hand, there is a strong extra-cellular matrix rich in collagen and secreted mainly by chondrocytes and early pillar cells across the filament {Morgan:1974fi}, adjacent to the place in which we characterized homeostatic stem cells.

**Figure 7.**
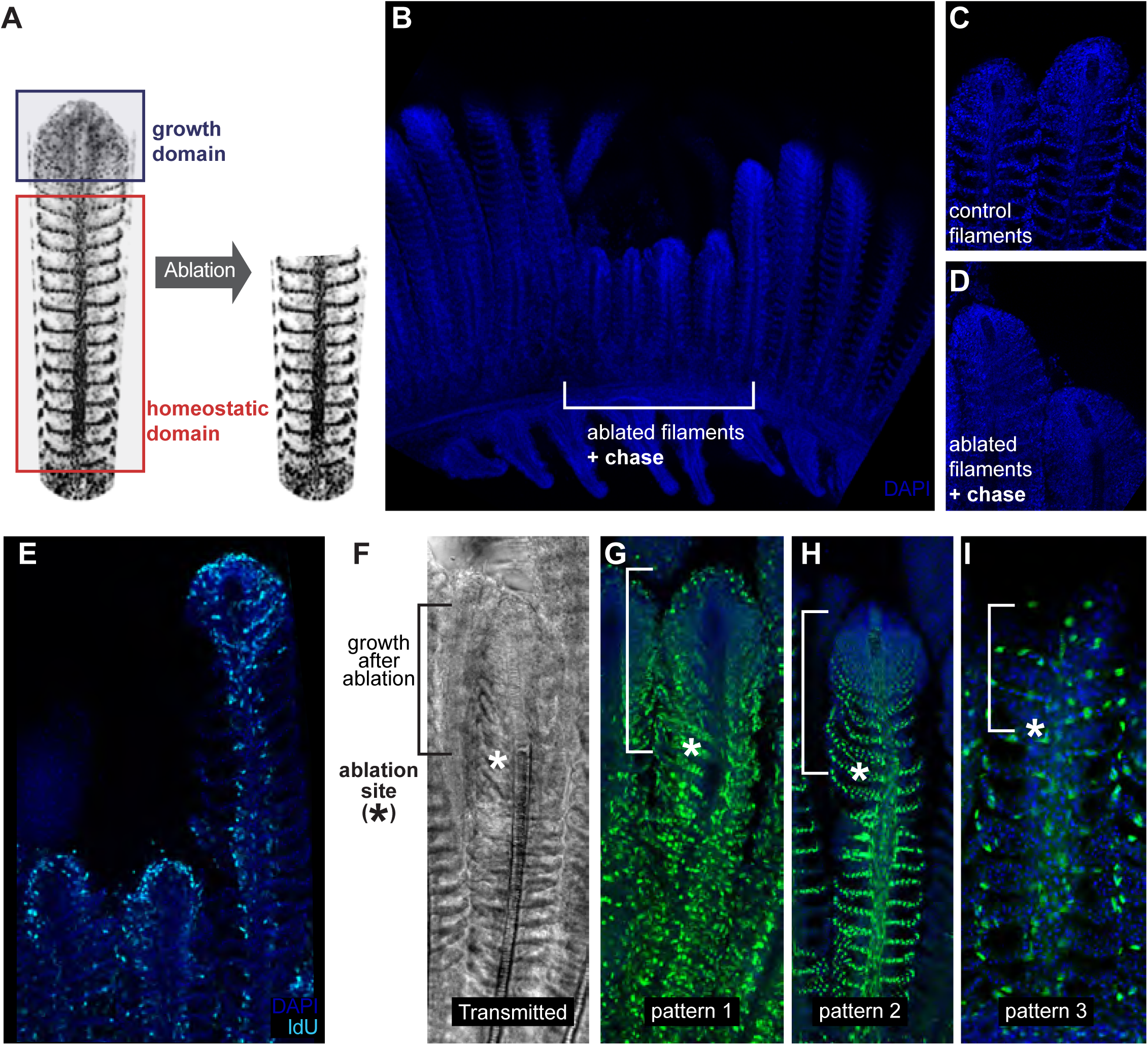
The Homeostatic Domain Sustains Growth After Filament Ablation. (**A**) Scheme of the ablation procedure. The growth domain and the upper part of the homeostatic domain are mechanically ablated. (**B**) DAPI image of control filaments shows an intact growth domain at the top. (**C**) DAPI image of injured filaments after a chase of one month shows a regenerated growth domain. (**D**) During the duration of the experiment, ablated filaments were unable to reach the length of their neighbour, non-ablated filaments

We speculated that modifying the close environment of homeostatic stem cells by ablating the growing zone of a filament could elicit a growth response from the homeostatic domain. We therefore ablated filaments by physically removing their upper region, where the growing domain and part of the homeostatic domain are located (Figure 7A). When experimental fish were grown for a month after ablation, we could still recognise the ablated filaments due to their shorter length, compared to that of their neighbour, non-ablated filaments (Figure 7B). Ablated filaments, however, restored the characteristic morphology of a growth domain at their most upper extreme (Figure 7C, D). Additionally, BrdU incorporation showed that the *new* growth domains were proliferative, showing a similar BrdU label than non-ablated filaments in the same branchial arch (Figure 7E-G). Our lineage analysis during homeostatic growth in medaka revealed different growth and homeostatic stem cells in each filament that maintained their fate during the entire life of the fish. We therefore wanted to assess whether the reconstitution of a filament growth domain after injury required cells from all different lineages or if alternatively, cells from a given lineage would change their fate to contribute to multiple recombination patterns (Figure 4 C-F). Injury paradigms have been shown to affect the fate commitment of stem cells in different models (Van Keymeulen *et al*, 2011; Suetsugu-Maki *et al*, 2012) while in others, proliferative cells maintain their fate during the regeneration process (Kragl *et al*, 2009; Knopf *et al*, 2011).

To address the nature of the cells re-establishing the growth domain, the same injury assay was performed on Gaudí^*Ubiq.iCRE*^ Gaudí^*RSG*^ transgenic fish that had been induced for sparse recombination at late embryonic stage (8 dpf) and grown for two months. When we analysed these samples 3 weeks after injury, we observed that the recombination pattern of the basal, non-injured region was identical to the recombination pattern of the newly generated zone (Figure 7F-I) (N=30 filaments in 6 branchial arches, N=17 for pattern 1, N=11 for pattern 2, N=2 for pattern 3). These results indicate that the re-established growth zone is formed by an ensemble of cells from the different lineages, and strongly suggest that homeostatic stem cells within all lineages can be converted to growth stem cells during regeneration. Our data definitively reveal that filaments possess the ability to resume growth from the homeostatic domain in a process that require cells from the different lineages. Overall, we propose from our observations that the different niches – physical and/or molecular - along the filament could operate as main regulators of the homeostatic-or-growth activity for stem cells in the fish gill.

## Discussion

In this study, we use mathematical modelling and genetic lineage analysis to reveal the rationale behind the permanent post-embryonic growth in a vertebrate. We introduce the fish gill, and particularly branchial arches, as a new model system that displays an exquisite temporal/spatial organisation, and use it to characterise growth and homeostatic stem cells. We reveal two domains harbouring growth stem cells: both extremes of each branchial arch contain *br-arch*SCs, which in turn generate *filam*SCs that locate to the tip of newly formed filaments. Additionally, *filam*SCs generate homeostatic stem cells at the lamellae along the longitudinal axis of the filament. The peripheral-to-central axis of branchial arches reflects a young-to-old filament order, and the longitudinal axis of a filament reflects a young-to-old lamellae order. The two growth stem cells and the one homeostatic stem cell types are clonal and organised in a hierarchical manner.

Our observations indicate that the relative position within the organ has a major impact on the growth vs homeostatic activity of stem cells. We have found that when the growth domain of a filament is lost, the homeostatic domain is able to generate a new, functional growth domain. This observation suggests that physical or molecular modifications in the local environment (relaxation of the inner core, or the absence of a repressive signal, respectively) could convert homeostatic stem cells into growth stem cells. In the absence of specific markers to label homeostatic stem cells before the ablation, however, we cannot discard the presence of quiescent stem cells that get activated after injury, nor the possibility of injury-triggered trans-differentiation as shown in the zebrafish caudal fin (Knopf *et al*, 2011).

Permanent post-embryonic growth is a challenging feature for an organism since new cells have to be incorporated to a functional organ without affecting its physiological activity. Restricting growth stem cells to the growing edge is an effective way to compartmentalise cell addition and organ function. Strikingly, the location of growth stem cells in gill filaments is highly reminiscent of the overall topology of meristems in plants (Greb & Lohmann, 2016). In both systems, axis extension occurs by the sustained activity of stem cells that locate to the growing edge. These stem cells consistently remain at the growing zone, while their progeny start differentiation programs and occupy a final location at the coordinates in which they were born. It is to note that other ever-growing organs in fish follow the same growing principle, with tissue stem cells located at the growing edge and differentiated progeny left behind, as it has been nicely shown for different cell types in the zebrafish caudal fin (Tu & Johnson, 2011) and the medaka neural retina and retinal epithelium (Centanin *et al*, 2011; 2014). Since stem cells are thought to have evolved independently in the vegetal and the animal lineages (Meyerowitz, 2002; Scheres, 2007), our results illustrate how the same rationale to sustain permanent growth can be adopted in the most diverse systems.

We have performed an organ-scale lineage analysis at cellular resolution and found that growth stem cells and homeostatic stem cells are fate restricted. We used two un-biased labelling approaches (ubiquitous expression of the inducible ErT2CRE and heat-shock induced expression of CRE) to identify at least four different fate-restrictions for gill stem cells, which generate reproducible labelling patterns along gill filaments. Since each filament contains all four fate restricted stem cells (we have not observed filaments lacking one entire lineage), our results determine that the growth zone of a gill filament is indeed an *ensemble* — a group of stem cells with different potencies that work in an interconnected manner. Two relevant avenues open from this analysis, namely: a) how stem cells are recruited together to a newly forming filament, a process that happens hundreds of times during the lifetime of a medaka fish and thousands of times in longer-lived teleost fish, and b) how stem cells coordinate their activity to maintain the ratio of cell types in the individual filaments. We have observed that the relative proportion of differentiated cells types is maintained along the filament axis, which once again points at a coordinated pace of cell type generation that is maintained life-long. One fundamental aspect to start addressing coordination is to define the number of stem cells for each lineage, a parameter that proved to be hard to estimate for most vertebrate organs. The prediction for gill filaments is that they contained a very reduced number of stem cells, for they generate all-or-none labelled filaments of a given cell type reflecting a clonal nature. Altogether, we believe that our results position the fish gill as an ideal system to quantitatively explore a stem cell niche hosting multiple lineage-restricted stem cells.

In most adult mammalian organs, stem cells maintain homeostasis by generating new cells that will replace those lost during physiological or pathological conditions. We have functionally identified homeostatic stem cells in the fish gill, and focussed on the ones generating pillar cells. Our lineage analysis demonstrates that growth and homeostatic stem cells are clonal along a filament, where the former generate the latter. The most obvious difference between these two stem cell types is their relative position; growth stem cells are located at the growing tip, beyond the rigid core that physically sustains the structure of the filament, while homeostatic stem cells are embedded inside the tissue, adjacent to the collagen-rich chondrocyte column. It is to note that both the function and the relative location of the gill homeostatic stem cells match those of the mammalian homeostatic stem cells, being located at a fixed position and displacing their progeny far away - as it is observed for intestinal stem cells, skin stem cells and oesophagus stem cells (Barker *et al*, 2008; Blanpain & Fuchs, 2009; Seery, 2002). The comparison of growth and homeostatic stem cells in the gill suggests the existence of a physical niche that would restrict stem cells to their homeostatic role, preventing them to drive growth. We believe that during vertebrate evolution, the transition from lower (ever-growing) to higher (size-fixed) vertebrates involved restraining the growth activity of adult stem cells. One of the main functions of mammalian physical niches, in this view, would be to restrict stem cells to their homeostatic function. Many stem cell-related pathological conditions in mammals involve changes in the microenvironment including physical aspects of the niche (Brabletz *et al*, 2001; Vermeulen *et al*, 2010; Ye *et al*, 2015; Oskarsson *et al*, 2011; Liu *et al*, 2012; Butcher *et al*, 2009), suggesting that homeostatic stem cells could drive growth in that context. Along the same line, the extensive work using organoids that are generated from adult homeostatic stem cells, like intestinal stem cells, (Sato *et al*, 2009; Kretzschmar & Clevers, 2016), demonstrates that healthy aSCs have indeed the capacity to drive growth under experimental conditions and when removed from their physiological niche. Our work, therefore, illustrates how different niches affect the functional output of clonal stem cells driving growth and homeostatic replacement in an intact *in vivo* model.

## Material & Methods

### Fish Stocks

Wild type and transgenic *Oryzias latipes* (medaka) stocks were maintained in a fish facility built according to the local animal welfare standards (Tierschutzgesetz §11, Abs. 1, Nr. 1). Animal handling and was performed in accordance with European Union animal welfare guidelines and with the approval from the Institutional Animal Care and Use Committees of the National Institute for Basic Biology, Japan. The Heidelberg facility is under the supervision of the local representative of the animal welfare agency. Fish were maintained in a constant recirculating system at 28°C with a 14 h light/10 h dark cycle (Tierschutzgesetz 111, Abs. 1, Nr. 1, Haltungserlaubnis AZ35–9185.64 and AZ35–9185.64/BH KIT). The wild type strain used in this study is Cab, a medaka Southern population strain. We used the following transgenic lines that belong to the Gaudí living toolkit (Centanin *et al*, 2014): Gaudí^*Ubiq.iCre*^, Gaudí^*Hsp70.A*^, Gaudí^*loxP.OUT*^ and Gaudí^*RSG*^.

### Generation of clones

Clones were generated as previously described (Centanin *et al*, 2014; 2011; Seleit *et al*, 2017; Rembold *et al*, 2006). A brief explanation follows for the different induction protocols. Fish that displayed high recombination were discarded for quantifications on lineage analysis and fate restriction to ensure clonality.

<u>Inducing recombination via *heat-shock*</u>: double transgenic Gaudí^*RSG*^, Gaudí^*Hsp70.A*^ embryos (stage 32 to stage 37) were heat-shocked using ERM at 42°C and transferred to 37°C for 1 to 3h.

<u>Inducing recombination via tamoxifen</u>: double transgenic Gaudí^*RSG*^, Gaudí^*Ubiq.iCre*^ fish (stage 36 to early juveniles) were placed in a 5µM Tamoxifen (T5648 Sigma) solution in ERM for 3 hours (short treatment) or 16 hours (long treatment), and rinsed in abundant fresh ERM before returning them to the plate. Adult fish were placed in a 1µM Tamoxifen solution in fish water for 4 hours, and washed extensively before returning them to the tank.

<u>Generating clones via blastula transplantation</u>: between 25 - 40 cells were transplanted from a Gaudí^*loxP.OUT*^ heterozygous to a wild type, unlabelled blastula. Transplanted embryos were kept in 1xERM supplemented with Penicillin-Streptomycin (Sigma, P0781, used 1/200) and screened for EGFP+ cells in the gills during late embryogenesis.

### Antibodies and staining protocol

For immunofluorescence stainings we used previously described protocols (Centanin et al., 2014). Primary antibodies used in this study were Rabbit a-GFP, Chicken a-GFP (Invitrogen, both 1/750), Rabbit a-Na^+^K^+^ATP-ase (Abcam ab76020, EP1845Y, 1/200) and mouse a-BrdU/Idu (Becton Dickinson, 1/50). Secondary antibodies were Alexa 488 a-Rabbit, Alexa Alexa 647 a-Rabbit, Alexa 488 a-Chicken (Invitrogen, all 1/500) and Cy5 a-mouse (Jackson, 1/500). DAPI was used in a final concentration of 5ug/l.

To stain gills, adult fish were sacrificed using a 2 mg/ml Tricaine solution (Sigma-Aldrich, A5040-25G) and fixed in 4% PFA/PTW for at least 2 hours. Entire Gills were enucleated and fixed overnight in 4% PFA/PTW at 4C, washed extensively with PTW and permeabilised using acetone (10-15 minutes at -20C). Staining was performed either on entire gills or on separated branchial arches. After staining, samples were transferred to Glycerol 50% and mounted between cover slides using a minimal spacer.

### BrdU or IdU treatment

Stage 41 juveniles were placed in a 0,4mg/ml BrdU or IdU solution (B5002 and I7125 respectively, Sigma) in ERM for 16 hours and rinsed in abundant fresh ERM before transferring to a tank. Adult fish were placed in a in a 0,4mg/ml BrdU or IdU solution in fish water for 24 or 48 hours, and washed extensively before returning them to the tank.

### Imaging

Big samples like entire gills or whole branchial arches were imaged under a fluorescent binocular (Olympus MVX10) coupled to a Leica DFC500 camera, or using a Nikon AZ100 scope coupled to a Nikon C1 confocal. Filaments were imaged mostly using confocal Leica TCS SPE, Leica TCS SP8 and Leica TCS SP5 II microscopes. When entire branchial arches were imaged with confocal microscopes, we use the Tile function of a Leica TCS SP8 or a Nikon C2. All image analysis was performed using standard Fiji software.

### Modelling

To model progenitor and stem cell scenarios for the addition of post-embryonic filaments we performed stochastic simulations for each considering a stretch of 6 filaments, and then compared them to experimental data. We chose stretches of 6 filaments because those guaranteed that we would be focussing on the post-embryonic domain of a branchial arch. A random filament would contain ca. 8 embryonic filaments, and we considered branchial arches with 20 or more filaments, which results in 6 post-embryonic filaments at each side.

<u>Stem cell model</u>: if there is only one stem cell in the niche, then all 6 filaments will share the same label, either 0 or 1. We draw random numbers from a Bernoulli distribution, where the probability parameter equals the experimental labelling efficiency of our dataset.

<u>Progenitor model</u>: in a similar manner, we considered the case of having 6 progenitor cells in the niche. Thus, this time a Bernoulli process of 6 trials with probability parameter equal to the labeling efficiency of the gill was simulated for each branchial arch.

<u>Experimental data</u>: We collected data from 22 Gaudí^*Ubiq.iCRE*^ Gaudí^*RSG*^ recombined gills, which we dissected and analysed under a confocal microscope and or macroscope - 8 to 16 branchial arches per gill. Subsequently, quantifications were done on the 6 most peripheral filaments from each side of a branchial arch. The labelling efficiency was estimated for each gill by employing a combinatorial approach: the number of labeled filaments at position +6 (i.e. oldest filaments selected) divided by the total number of branchial arches analysed for that gill.

<u>Comparison:</u> To compare each model to the experimental data, we compute an objective function in the form of a sum of square differences for each gill and each model. The smaller this objective function is, the better the fit between experimental data and simulations. We annotated both the number of switches and of labelled filaments in each branchial arch.

There exist 19 possible pairs *(s,f)* of switches and labelled filaments, ranging from (0,0), (0,6) up to (5,3). We calculated for each pair *i*, of the form *(s,f)* the frequency of observing it in the data from each gill *j*, 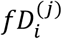, and in simulations of 5000 filament stretches per gill *j*, 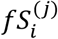. The objective function *f*^(*j*)^ was computed for each gill as an adjusted sum of square differences:

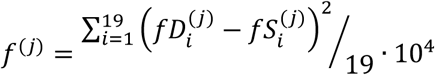

This was done for both the stem cell and the progenitors models. The factor 10^4^ was introduced for avoiding small numbers thus facilitating the comparison between results. The procedure was repeated 1000 times, producing 1000 objective functions per gill and per model, and therefore obtaining an average value and a standard deviation for each gill for each model.

## Acknowledgements

We thank S. Lemke, J. Wittbrodt, J. Lohmann and G. Begeman for scientific inputs at earlier versions of this project, and A. Seleit, K. Gross and I. Krämer for active discussions and suggestions on the manuscript. We are grateful to U. Engel and the Nikon Imaging Center for advice and support with microscopes and imaging, and to E Leist, A Sarraceno and M Majewski for fish maintenance. This work has been funded by the Deutsche Forschungsgemeinschaft (German Research Foundation, DFG) via the Collaborative Research Centre SFB873 (subproject A11 to LC and B08 to AMC). JS is the recipient of a Melbourne Research Scholarship from the University of Melbourne, Australia.

## Author Contributions

JS & EMA conducted most experiments, analysed the data and edited the manuscript, D-PD & AMC run mathematical simulations and models and analysed *in silico* and experimental data, DAE provided support and hosted JS, KN performed experiments and provided reagents, LC conceived the project, performed experiments, analysed the data and wrote the manuscript with support from JS, EMA, D-PD & AMC.

